# Fast neutron mutagenesis in soybean creates frameshift mutations

**DOI:** 10.1101/2020.11.17.387373

**Authors:** Skylar R. Wyant, M. Fernanda Rodriguez, Corey K. Carter, Wayne A. Parrott, Scott A. Jackson, Robert M. Stupar, Peter L. Morrell

## Abstract

The mutagenic effects of ionizing radiation have been used for decades to create novel variants in experimental populations. Fast neutron (FN) bombardment as a mutagen has been especially widespread in plants, with extensive reports describing the induction of large structural variants, i.e., deletions, insertions, inversions, and translocations. However, the full spectrum of FN-induced mutations is poorly understood. We contrast small insertions and deletions (indels) observed in 27 soybean lines subject to FN irradiation with the standing indels identified in 107 diverse soybean lines. We use the same populations to contrast the nature and context (bases flanking a nucleotide change) of single nucleotide variants. The accumulation of new single nucleotide changes in FN lines is marginally higher than expected based on spontaneous mutation. In FN treated lines and in standing variation, C→T transitions and the corresponding reverse complement G→A transitions are the most abundant and occur most frequently in a CpG local context. These data indicate that most SNPs identified in FN lines are likely derived from spontaneous de novo processes in generations following mutagenesis, rather than from the FN irradiation mutagen. However, small indels in FN lines differ from standing variants. Short insertions, from 1 – 6 base pairs, are less abundant than in standing variation. Short deletions are more abundant and prone to induce frameshift mutations that should disrupt the structure and function of encoded proteins. These findings indicate that FN irradiation generates numerous small indels, increasing the abundance of loss of function mutations that will impact single genes.

**Significance Statement:** Irradiation mutagenesis is commonly viewed as a method to induce large structural variants in genomes. We also find enrichment in small insertion and deletion (indel) variants. The radiation-mutagenized lines averaged 32 indels per line, far exceeding the number estimated to occur by spontaneous processes, indicating that these arose from the irradiation treatment. Nevertheless, naturally-occurring standing variation among soybean accessions is still four orders of magnitude higher than the level of diversity introduced by mutagenesis. Induced mutations from any source are likely to constitute a relatively small portion of the genetic variation present in crop species. However, irradiation mutagenesis is useful for altering genomes by introducing small indels into single genes or disrupting gene clusters by creating structural variants.

## Introduction

Naturally occurring mutations have long been recognized as the primary source of genetic variation used for selection in plant breeding programs. Mutations induced by irradiation and mutagenic chemicals have also been important for generating variation when naturally occurring genetic variation for a trait was absent or insufficient (1). The advent of CRISPR and other site-directed nucleases has enabled targeted nucleotide changes, thus eliminating much of the randomness from the generation of de novo variation. Nevertheless, not all genomic sites lend themselves to editing for reasons that are still not clear (2).

The advent of genetic engineering technology for crops cast mutations in a new light. The US Food and Drug Administration initially speculated that unintended mutations caused by the insertion of a transgene or by mutations occurring during the tissue culture portion of the transformation process could trigger the production of unknown toxins, negatively affecting human food and animal feed safety (3). As a result, an extensive and expensive system for testing the safety of genetically modified crops has been instituted (4, 5). It was also hypothesized that these mutations could cause crops to have undesirable environmental effects (6, 7). More recently, the potential for CRISPR and other site-directed nucleases to induce genetic changes beyond targeted nucleotide sites (i.e., “off-targets”) has created renewed interest in genome-level screens for de novo mutations, in part due to lingering concerns over the safety implications of these off-target effects (8, 9). The best way to assess the safety implications of off-target edits is by comparison to natural and induced mutations that have a history of safe use in breeding programs.

Comparing naturally occurring variants to variants in mutagenized lines provides a best-case scenario for determining how and if induced mutations differ from naturally occurring variants. Mutagenized plants with heritable phenotypic differences from the original experimental line must carry mutations capable of altering the phenotype. Mutagenesis experiments use a mutagen exposure at a level that would be lethal to a proportion of treated individuals. Standard measures of toxicity use an “LD_50_” (median lethal dose), meaning that half of all treated individuals do not survive the mutagenic treatment at the level of application. Mutagenized individuals that survive are likely to have been subject to mutation but carry induced genetic changes that are less damaging.

While the mutagenic effects of FN radiation have been successfully applied to create genetic novelty in plants, (10), the full spectrum of FN-induced mutations is poorly understood. FN bombardment can create large structural variants, including deletions, duplications, inversions, and translocations, but single nucleotide point mutations and small indels have also been reported (11). Previous studies have characterized large deletions and duplications in soybean (12, 13). In *Arabidopsis thaliana*, there was enrichment for single-base substitutions, primarily at pyrimidine dinucleotides (11). In rice, single-base substitutions constituted the most abundant mutation type, but deletions mutated the largest number of genes (14).

Distinct patterns of mutation tend to predominate for both spontaneous and mutagen-induced changes. For spontaneous mutations, transitions (changes between pyrimidines or between purines) are observed at higher relative rate than transversions (changes between pyrimidines and purines) (15). New mutations can be quite distinct from standing variation in populations, a phenomenon explored in deep-resequencing panels (16). Short insertions and deletions, typically smaller than a sequence read length, are also readily detectable with Illumina short-read sequencing (17). Short deletions relative to a reference genome are potentially the most readily detectable indel events, owing to fewer issues with initial read mapping. Short indels may be particularly abundant in low complexity sequence, including mononucleotide repeats. Low complexity sequence also provides challenges due to limitations of sequencing chemistry and alignment algorithms.

Mutation rates vary across the genome, though the causes of this variation are not completely understood (18). Chromosomal-scale patterns in mutation rate are impacted by factors such as GC-content (16, 19), local recombination rates (16), and methylation rates (20). However, the immediate context of variants appears to have the greatest impact on the nature of variants observed (21), see also (22, 23). The most abundant mutation observed in many organisms is the C→T transition, frequently in a CpG context (24, 25). That is, a cytosine is bound by a phosphate bond to guanine on the same side of the nucleotide strand (or nucleoside) and in the 5’ to 3’ orientation. As a shorthand for mutations that could have arisen on either strand, we use C→T*, which refers both to C→T changes and the reverse complement G→A. These changes cannot typically be distinguished in resequencing studies. The effect of context varies based on mutagenizing factors. For example, the mutagen ethyl methanesulfonate, when used in plants, tends to induce transitions in a GC-rich context (26). These results suggest that observed mutations and the context in which they occur are mutagen-dependent and based on specific biochemistry. The specific context in which mutations are more likely to occur matters because the immediate flanking nucleotides can make certain classes of large-effect mutations, such as stop codons, less probable.

Quantification of the influence of local nucleotide sequence context on mutation requires that mutations are divided into specific classes. Identification of the mutational context of insertions and deletions is often not directly observable because multiple equiprobable local nucleotide sequence alignments are possible around the mutation (27, 28). There are 12 possible single-nucleotide mutations. If the local context affects the potential occurrence of these variants, particular nucleotides would be over-represented among the immediately flanking sequences. Ideally, an approach would also account for the probability of sampling each of the nucleotides rather than relying on an equiprobable occurrence of all four nucleotides. It should also permit the examination of first or second-order interactions in terms of the presence of particular flanking nucleotides (29).

Here we investigate the differing nature of induced versus standing variants among panels of different soybean genotypes. This includes 27 soybean lines subject to FN mutagenesis compared to standing variation in 107 diverse soybeans. Because the potential for creating large structural changes is well documented (12), we focus on single nucleotide, and small indel variants that can be readily detected with short sequence reads (30). We start by determining if there is a difference in the mutational spectrum of induced mutations relative to standing variation. We then investigate if local context (flanking nucleotides) affects the frequency and type of mutations.

## Results

### Identification of Standing and De Novo Variants

To examine the effects of FN mutagenesis on single nucleotide variation in soybean, we made use of two published resequencing data sets (12, 31) that included 27 soybean lines subject to 8, 16, or 32 gray units (Gy) of FN radiation (Table S1, Figure 1). We also examined one non-mutagenized soybean inbred line (M92-220) that served as the initial parental line for the FN mutant population. Sequenced accessions were subject to four to eight generations of self-fertilization before sequencing (Table S1).

**Fig. 1.**
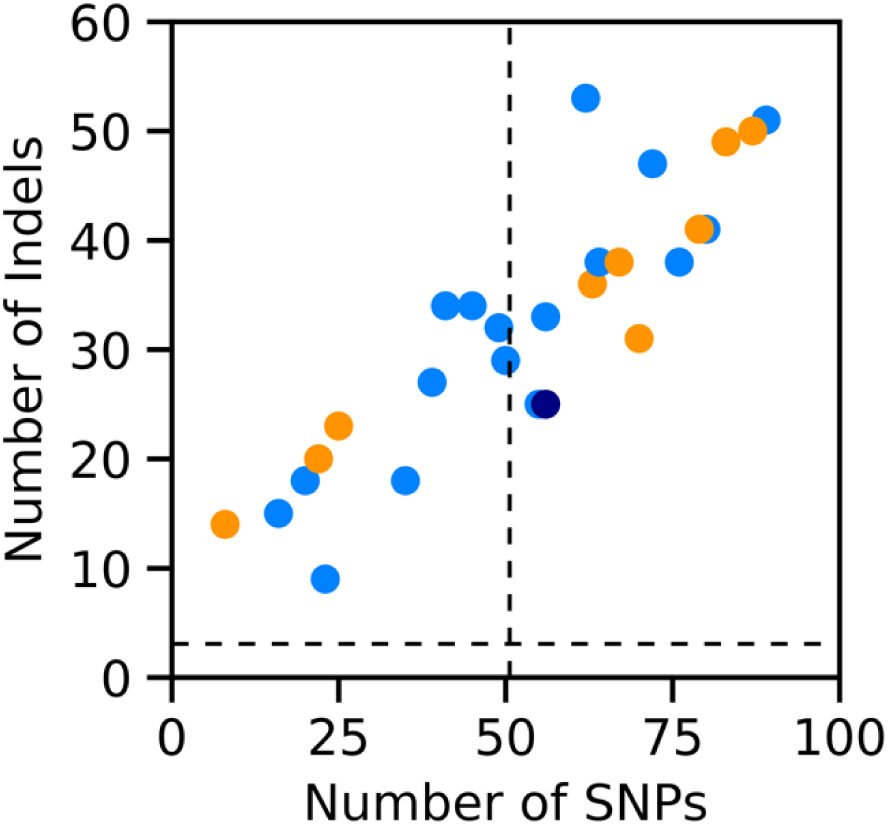
The number of variants identified in each FN line. The samples were exposed to 8 Gy (dark blue), 16 Gy (light blue), or 32 Gy (orange). The expected number of SNPs and indels based on average mutation rates are indicated by dotted lines.

Our goal was to identify induced mutations in the M92-220 treatment lines subject to FN exposure. A necessary first step was the identification of variants that differentiate M92-220 from the ‘Williams 82’ reference genome (32). To identify these variants, we included resequencing from prior studies (12, 31), along with newly collected Illumina resequencing from M92-220 with and without 10X Genomics linked reads. To provide a contrast between induced mutations found in treatment lines and natural variation, we made use of a published resequencing dataset of 106 soybean lines reported by (33). The 106 line dataset (33) was also used to filter naturally-occurring variants from our FN-treated lines.

Reference-based read mapping from three individuals of the M92-220 inbred line identified a total of 1,668,049 single nucleotide variants with an observed transition/transversion ratio (Ts/Tv) of 1.90. We also identified 299,980 sequence insertions or deletions (indels) smaller than 280 base pairs (bp), which are differences from the Williams 82 reference (Supplemental Data S1). 10X Genomics linked reads provided improved potential to detect intermediate-length structural variants. Using a single accession of M92-220 sequenced with 10X Genomics linked reads, we identified 495 additional indels that were not identified by paired-end sequencing alone, ranging in size from 17,767 to 499,355 bp. Indels include 105 deletions relative to the reference, 147 duplications, 16 inversions, ten translocations, and 217 variants with types that could not be identified. Deletions in M92-220 relative to the Williams 82 reference constituted 3.82% of the total genome size. De novo variants could not be detected in regions that harbored deletions relative to the reference, so these regions were masked from further analysis.

Standing variation, in terms of reference-based read mapping of the 106 samples from (33) and M92-220 from (12, 31) resulted in the identification of 9.7 million SNPs with an observed Ts/Tv = 1.94 and 1.5 million indels, for a total of 11,196,099 variants (Supplemental Data S2). Estimated pairwise diversity (34), here reported as θ_π_ = 4N_e_µ at all SNP sites, is shown for chromosome 1 in Figure S1A. Genome-wide diversity was θ_π_ ∼2 x 10-3 in the full panel from (33), which included soybean cultivars, landraces, and wild accessions that represent the variation present in the modern varieties or that is accessible to breeders.

Standing variants were partitioned into two classes based on frequency to allow for more direct comparison with FN variants. The first class included common variants with a non-reference allele count of at least three in our dataset. The second class included rare variants with a non-reference allele identified in either one or two genotypes. These rare variants have experienced fewer generations of selection and are likely more similar in terms of mutational spectrum and context to variants produced by spontaneous mutational processes.

### Variation in FN Lines

FN lines exhibited a total of 2,296 de novo variants in the homozygous state, including indels, for a mean of 85.3 (± 34.0) new mutations per line. This total included variants distributed relatively evenly across all 20 soybean chromosomes and 24 of the 1,170 scaffolds in the Gmax_275_v2.0 genome assembly (32). We identified 1,430 SNPs, which is an average of 53.0 (± 23.0) per line. We also identified 866 indels for an average of 32.1 (± 11.90) per line. Among these variants, those most likely to disrupt genes include 34 frameshift and 68 missense changes that impact an average of 2.15 (± 1.76) genes per plant.

We used observed variants across the 966 Mb reference genome in a haploid sample to estimate SNP-level diversity. This results in a pairwise diversity estimate of θ_π_ = 1.1 x 10-7 for de novo mutations (Figure S1B). Note that θ_π_ estimates are parametric and not dependent on sample size (34). Thus, the very low polymorphism observed in Figure S1B results from de novo mutations either from the FN treatment or due to spontaneous mutations arising during line maintenance.

Lines exposed to a higher dosage of fast neutron radiation (32 Gy) had a larger average number of SNPs and small indels than lines with a lower dosage (8 or 16 Gy) (Figure 1). However, there is a relatively large variance in mutation number, with the lowest number of observed variants occurring in an FN line with the highest (32 Gy) exposure (Figure 1). Another factor affecting observed diversity is the number of generations of maintenance of each line. After mutagenesis, lines were subject to four to eight generations of self-fertilization (Table S1). A more difficult factor to account for is the within-line heterogeneity present within the seed stock of M92-220 used for initial mutagenesis (35). For this analysis, we estimated that ten generations of self-fertilization (accompanied by lineage-independent spontaneous mutations) occurred before the mutagenesis treatment. Our observed number of SNP and indels per line are not well predicted by either the number of generations of line maintenance or dosage during a single generation of exposure to FN radiation. This is likely the result of a limited number of observations of a stochastic process.

We can estimate the expected number of mutations in the FN lines in the absence of the single-generation FN treatment based on average per generation mutation rates. We make use of the mutation rate estimates of (36) from a mutation accumulation experiment in *Arabidopsis thaliana*. Using the mean (6.53 x 10^−9^) of nucleotide substitution rates estimated by (36), we expect a mean of 50.57 SNPs per line. Using a mean indel mutation rate estimate of 0.47 x .10^−9^ (36), we expect a mean of 3.08 indels per line (Figure 1). The observed number of SNPs per line is very close to the expectations based only on spontaneous mutations. The observed number of indels per line is more than an order of magnitude greater than expected based on spontaneous variants over generations.

### Mutational Spectrum

The SNP mutational spectrum is similar for the 107 diverse lines and the 27 FN lines (Figure 2). Transitions are more common than transversions, with C→T* transitions being particularly abundant (Figure 2). Among the 107 lines, the second most abundant class of mutations is A→G* transitions (Figure 2), with common standing variants particularly enriched for this class. For FN lines, C→T* transitions are slightly more abundant than in the 107 lines in the standing variation panel, while A→G* transitions are slightly less abundant, as described below.

**Fig. 2.**
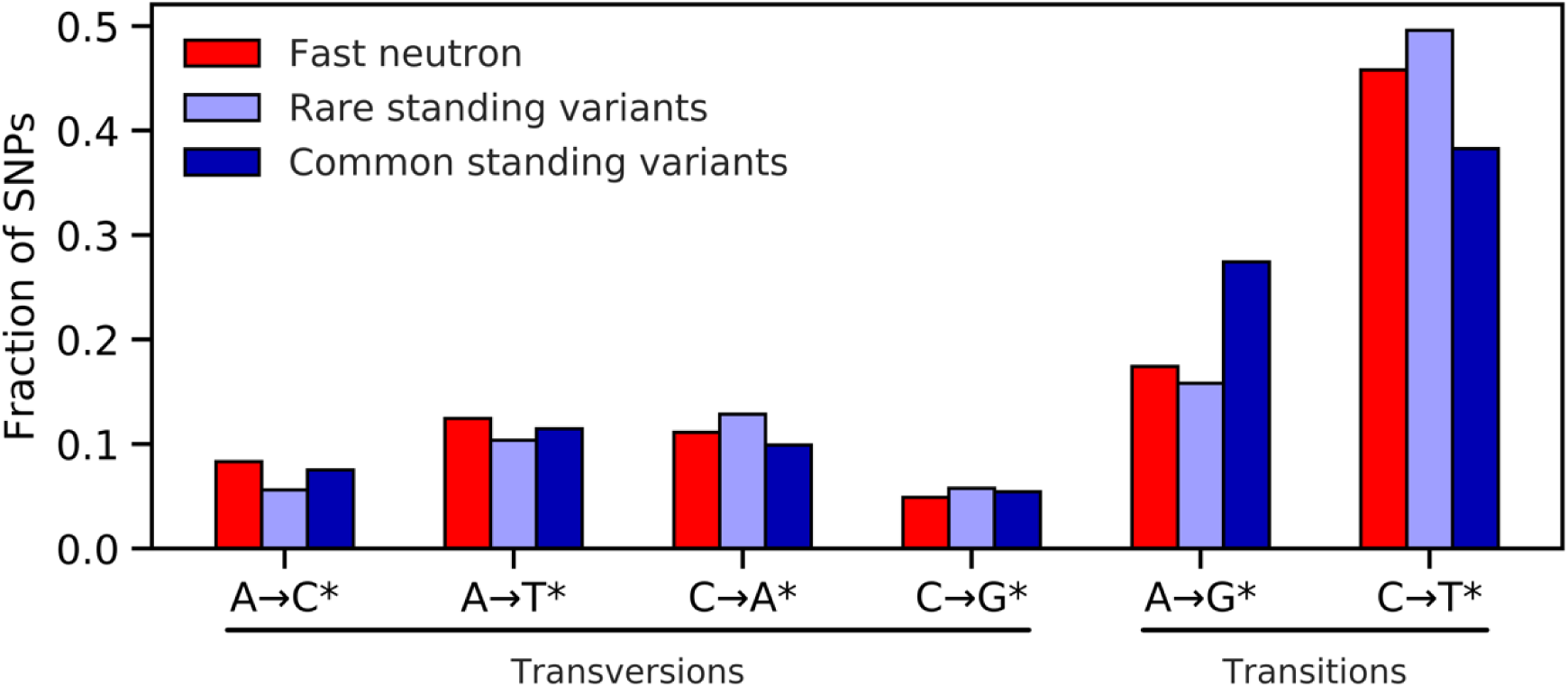
The mutational spectrum of FN variants, rare standing variants (non-reference allele count of two or fewer), and common standing variants (non-reference allele count of three or higher). Each class combines the displayed change and its reverse complement (e.g., the notation C→T* indicates that both cytosine to thymine and the reverse complement guanine to adenine mutations are binned into a single class (see text)).

### Context of Mutations

The survey of standing variation in the 107 lines identified 9,708,345 SNPs. The most abundant class of variants includes 4,035,491 (41.57%) C→T* transitions, and the least abundant includes 532,675 C→G* (5.49%) transversions. The very large sample size for standing variation provides the potential to observe all four nucleotides at each of the four flanking nucleotide positions. In standing variation, C→T mutations tend to be followed by G (guanine) at either position 1 or position 2 following the SNP (see Figure 3 for explanation). While the CG motif provides the context for the most abundant class of mutations, it is the rarest of all two nucleotide motifs in the soybean genome, constituting only 1.6% of all two nucleotide combinations. In contrast, the AA and TT motifs make up 11.9% of all two nucleotide combinations.

**Fig. 3.**
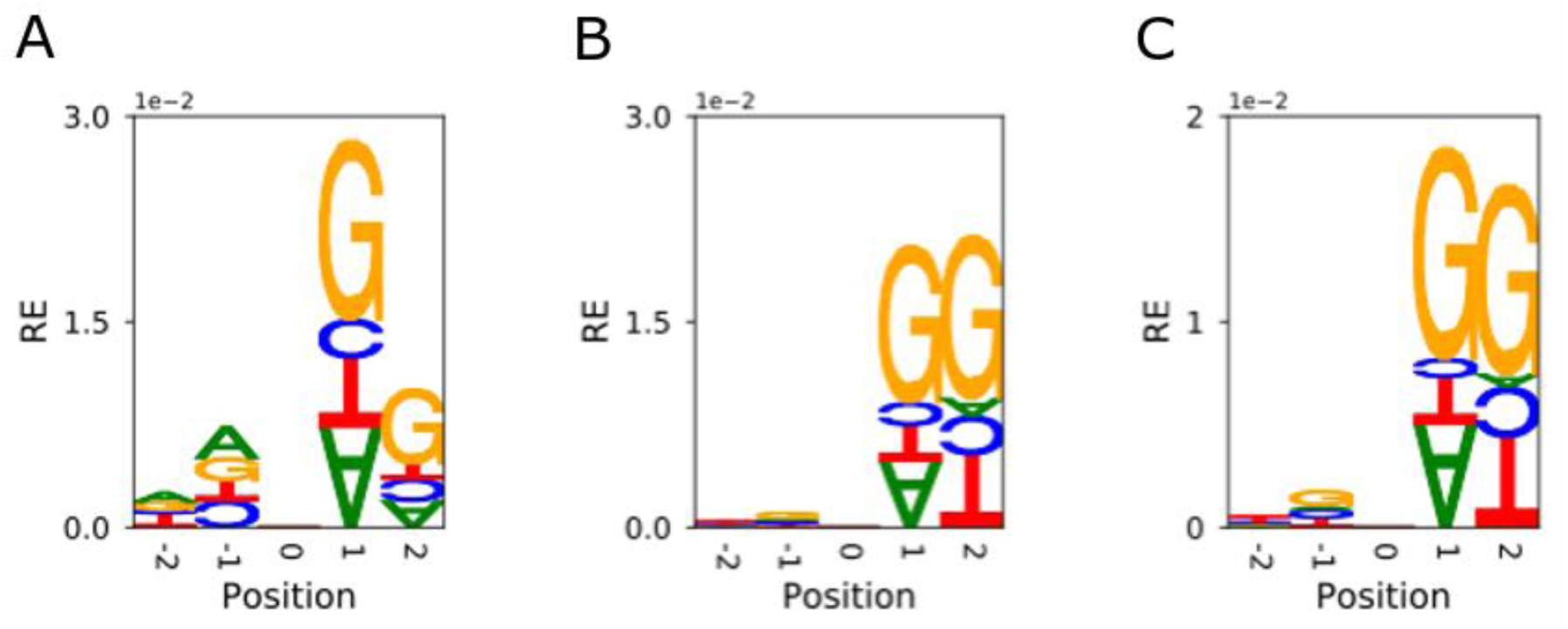
The nucleotide sequence context in which C→T transitions occur relative to the reference genome for FN variants (panel A, N = 319), rare standing variants (panel B, N = 647,614), and common standing variants (panel C, N = 1,350,269). The relative height of a letter indicates its relative entropy (RE), with overrepresented and underrepresented bases portrayed as right-side-up and upside down, respectively. The null expectation (zero relative entropy) is based on a nearby nucleotide of the same base that has mutated, chosen at random.

The contextual pattern for C→T transitions in the FN lines is similar to those of the diverse panel. However, comparison in de novo variants is limited by the number of observations, with C→T* constituting 655 of the 1,430 (45.8%) observed de novo variants. C→T changes in the FN lines were also found to be followed by G at positions 1 or 2, but the relative effect of position 1 appears to be larger (Figure 3B). The −1 position was typically A or G, but this was a smaller effect. However, the number of these combinations was limited by sample size.

### Indel Variation

The distribution of indel sizes differed between variants observed in the FN, rare standing, and common standing classes (Figure 4). In FN lines, there are many more deletions than insertions, with only 28.5% of indels as insertions. In standing variation. In standing variation, the two classes are more evenly represented, with 860,649 deletions and 756,095 insertions (46.8% insertions). Rare standing variants should be impacted less by purifying selection than the common standing variants, and thus are most similar to expectations for new mutations. Comparing the rarest indels in the standing variation to the same size class in the FN lines using a paired t-test identified significant differences for insertions (*p* = 0.0005) and for deletions (*p* = 0.008). For all classes of variants, short indels, especially single base pair indels, are the most abundant (Figure 4). Figure 4 depicts indels up to 20 base pairs, though observed variants include indels of up to 279 bp in length (Supplemental Data S1 and S2).

**Fig. 4.**
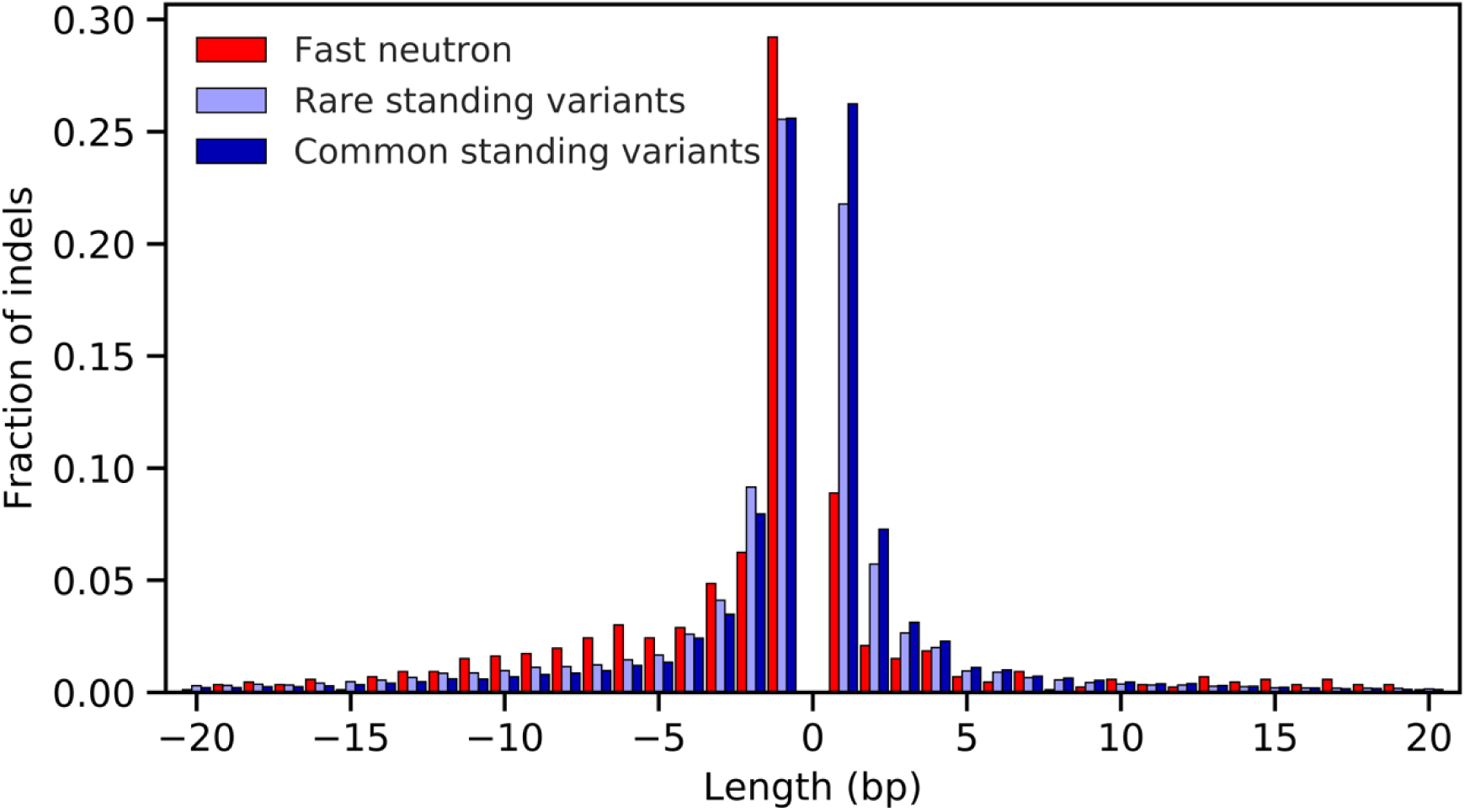
The distribution of insertion and deletion lengths of FN variants, rare standing variants (non-reference allele count of two or fewer), and common standing variants (non-reference allele count of three or higher). Insertions are shown as positive values and deletions as negative values. Variants with lengths greater than 20 bp are not shown.

Observed short indels in FN lines include 80 located within exons, with 34 of these changes introducing frameshifts that alter the amino acid sequence of a peptide. Based on Variant Effect Predictor (VeP) (37) analysis of all variants in both datasets, 26% of all coding variants in FN lines induce a frameshift. This compares to 3.9% of rare standing coding variation and 2.8% of common standing coding variation. The much more frequent observation of the disruption of coding genes among variants in FN lines maintained under single seed descent reflects the limited opportunity for purifying selection to act on de novo mutations.

Single base pair indels are more clearly identifiable as a single mutational event (38), and thus the nature of the nucleotide change is readily identified. The mutations in FN lines and standing variants show a similar tendency toward deletions of A or T. Previous studies in soybean have reported observed GC content of 33% (39), similar to the 34% observed here (Figure S2A). Single base pair insertions are even more heavily dominated by A and T changes (Figure S2B).

## Discussion

We examined levels of nucleotide sequence diversity in 27 soybean lines subjected to FN mutagenesis. Most assays of genetic variants include variants arising from a mixture of mutagenic processes (40). The primary difficulty is distinguishing FN-induced single nucleotide variants from those that arise from spontaneous mutations. The nature of mutations, their rate of occurrence, and the nucleotide sequence context in which variants are observed can be used to assess the proportion of soybean variants that were caused by FN mutagenesis as opposed to spontaneous processes. The observed SNP variants in the FN lines are primarily C→T*, which is the most abundant class of variants in soybean generally and occur at a rate that is consistent with variants expected to arise spontaneously in the seed stock or generations of inbreeding following mutagenesis. An examination of *Arabidopsis thaliana* lines treated with 60 Gy of fast neutron irradiation identified a ∼50-fold single generation increase in mutation rate (11). The same study noted an increase in G →T transversions that is not observed in our soybean lines. Aside from the difference in species examined, the highest level of FN exposure in the present study was 30 Gy, which may account for some of the differences in observed patterns of mutation. The lack of a distinctive pattern of mutation is in contrast to results previously published on the effects of chemical mutagens. This includes sodium azide in barley, where A→G* transitions predominate (41, 42) or ethyl methanesulfonate, which can result in alkylation of guanine, creating primarily C→T* transitions (26).

The most distinctive aspect of FN mutagenesis found in the present study is the induction of a large number of small deletions. This result closely aligns with the observations of (11), where the 253 single base pair deletions were the most abundant class of indel variants. Among the 34 frameshift that will oftentimes alter gene function, 18 are single-base deletions. FN-induced indels include proportionally fewer short 1-3 based pair insertions. The induction of small deletions that eliminate gene function is highly desirable for phenotypic screens aimed at identifying the genetic basis of observable phenotypes through gene knockdown or knockout. Of course, the primary challenge with FN mutagenesis is that double-stranded DNA breaks can create large structural variants, including deletions, which impact many genes, posing a challenge to associate phenotypes with individual mutations (43).

Efforts to identify induced nucleotide sequence changes must actively address several analytical and experimental issues. The relative magnitude of these issues, as measured by the number of variants they contribute, includes the following: First, experimental contamination through “off-type lines” or lines subject to unintended hybridization (or outcrossing) (26) can contribute millions of variants. Second, for reference-based read mapping, the differences between the line being investigated, and the reference genome must be identified and eliminated from comparisons (44). Third, levels of expected heterozygosity among induced variants must be considered, and sequencing depth must be sufficient for identification of heterozygous variants. Most experimental designs in plants involve inbred mutagenized lines. The number of generations of inbreeding subsequent to mutagenesis must be tracked to identify variants and to accurately estimate expected heterozygosity at induced mutations (26). Fourth, heterogeneity in experimental lines (35, 45) subject to the initial treatment can contribute large numbers of variants that continue to segregate or occur as fixed differences among treatment lines (44). Fifth, sequencing errors can contribute to low-quality variant calls (46). The sources of these errors include base call errors, inadequate coverage, poor mapping quality (Figure 5), repetitive DNA sequences, paralogous loci where reads cannot map uniquely, and quality issues in a reference genome (47).

**Fig. 5.**
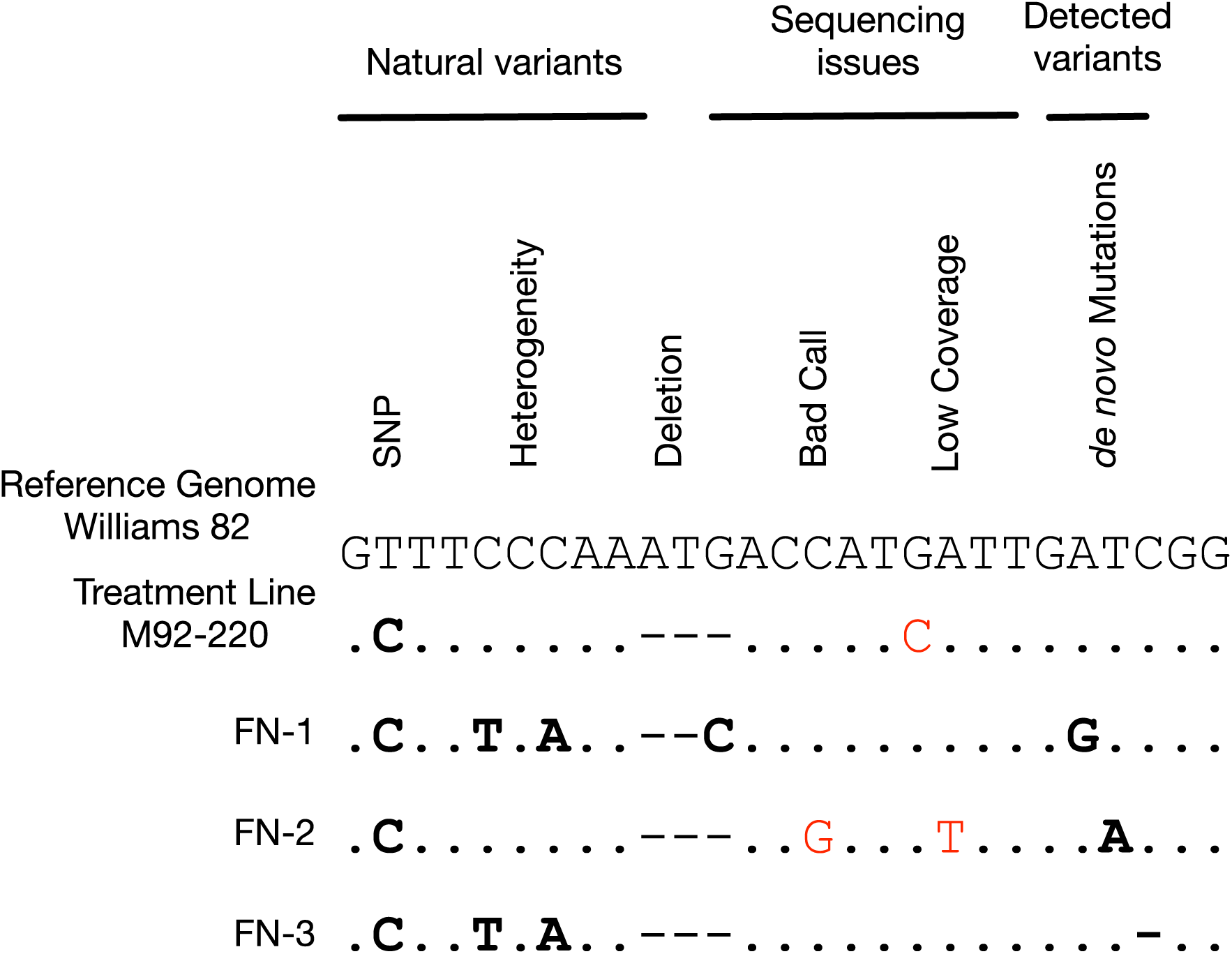
The detection of de novo mutations in FN treatment lines requires the filtering of naturally-occurring variants that are present in the initial treatment lines, including deletions in the treatment line where reference-based read mapping cannot identify variants. Sequencing issues, including bad variant calls and low coverage regions, are removed to isolate variants that are unique to treated lines. Base calls that are accurate are shown in bold, even if they are filtered out. Erroneous base calls are shown in red.

From a biological perspective, we find that the SNP frequency and sequence context of variants in the FN lines are similar to expectations based on the spontaneous rate of mutation over the generations of the experiment. This finding suggests that a large portion of single-nucleotide variants in the FN lines arise as spontaneous mutations, rather than as a direct result of the FN mutagenic process. Meanwhile, we find that FN mutagenesis induces many short indels, a portion of which result in frameshift mutations within the coding portions of genes. FN mutagenesis has long been exploited for experiments seeking to knock down or knock out single genes and to eliminate dominant traits in breeding programs; short indel-induced frameshift mutations are likely contributors.

From a regulatory point, these findings form a baseline with which to evaluate the relevance and importance of other mutations that result from the application of biotechnology. Applications of targeted genetic modifications in plants are rapidly expanding (48). Although the quantification of off-target effects from various genome editing applications is confounded by differences in methodology, the numbers reported are low, perhaps to the point of reflecting the background mutation rate. In contrast, technologies long considered safe can be quite disruptive at the whole-genome level.

## Materials and Methods

### Resequencing Data

To examine the effects of fast neutron (FN) mutagenesis on single nucleotide variation in soybean, we used published resequencing data sets that included FN-treated lines (12, 31) and untreated lines that serve as a control (33). Resequencing data from 30 lines subject to FN mutagenesis, all from the M92-220 line (12), were the treatment group. We also reanalyzed the sequence from three individuals from the M92-220 line that were non-mutagenized (Table S1). Soybean lines analyzed here were exposed to 8, 16, or 32 Gy of radiation and four to eight generations of self-fertilization (Table S1). Three of the 30 mutagenized lines were removed from analysis due to contamination detected during the de novo variant analysis. Of the remaining 27 lines, two sets of three siblings were derived from the same mutagenized plant (Table S1). None of the lines exposed to mutagenesis displayed aberrant phenotypes. We used published resequencing data from 106 soybean lines, including wild, landrace, and elite accessions reported by (33) and a single accession of the cultivated line M92-220 reported by (31) to identify naturally occurring variants in soybean.

A important step in the identification of de novo variants is the identifications of differences between the treated line and the reference genome and also heterogeneity and heterozygosity in the treatment line. We made use of previous Illumina paired-end resequencing of the original variety used for mutagenesis, M92-220. This included 100 bp Illumina sequencing from (31). We also collected additional whole-genome sequencing data from two different plants from the same seed lot, as there is known to be heterogeneity within the M92-220 seed stock (31). These newly collected sequences have been deposited in the NCBI SRA database, project number PRJNA670564. Reference-based read mapping is as described below, except for the 10X Genomics samples, which made use of the Longranger software version 2.2.2 https://support.10xgenomics.com/genome-exome/software/downloads/latest). The Longranger software was also used to identify structural differences between a reference genome and a query sample that are supported by both linked reads and split read information.

### Read Alignment and Variant Calling

Read alignment, and variant calling were implemented using publicly available software integrated with bash scripts in the ‘sequence_handling’ workflow (49). Configuration files and scripts used for analysis are available at https://github.com/MorrellLAB/Context_Variants_Soy. Reads were aligned to soybean reference genome “Gmax_275_v2.0,” a part of Phytozome 11 release (https://jgi.doe.gov/data-and-tools/phytozome/). Alignment parameters were adjusted to account for the levels of nucleotide diversity in soybean reported by (50).

The variant call format (VCF) (51) files for the 27 FN lines, 107 diverse lines, and four M92-220 lines were filtered with bcftools (52) to include only variants and sample genotypes which satisfied a set of quality criteria. Filtering criteria are implemented in a shell script in the project repository https://github.com/MorrellLAB/Context_Variants_Soy. Specifically, we filtered standing variants into two classes based on non-reference allele count. The common class contained all variants with an allele count of three or higher, and the rare class contained all variants with an allele count of two or less. VCF and related files are available through a Data Repository of University of Minnesota (DRUM) archive; available for review at: https://drive.google.com/drive/folders/1UsA9IKJRo25p8X1DIRAFMybLwtcmAtOb.

Descriptive statistics from VCF files, including observed heterozygosity and average pairwise diversity, were calculated using scikit-allel (53). The transition/transversion ratio, nature of variants relative to the reference (mutational spectrum), and SNP quality scores were calculated using bcftools (52).

### Identifying de novo variants

A major challenge with the identification of de novo variants is distinguishing variants present in a line prior to modification from de novo mutations. In practice, this involves filtering out several classes of variants, including; a) differences between the genotype used for mutagenesis and the reference genome, b) regions of the genome where variants cannot be identified, such as deletions in the mutagenized germplasm (that are a natural part of standing variation) relative to the reference, and c) heterozygosity within lines or heterogeneity among lines of the mutagenized germplasm (13, 44) (Figure 5). It is also necessary to account for variants that arise during seed maintenance prior to mutagenesis. Even if all lines are derived through single seed descent, a large seed increase over multiple generations is necessary to create the bulk seed subject to FN treatment. To account for this, we add ten generations of mutation to estimates of variation in the M92-220 seed lot before FN treatment.

Nucleotide sequence diversity in the FN panel was estimated using pylibseq, a Python interface for the libsequence library for evolutionary genetic analysis (54). All high-confidence SNPs in FN-lines are homozygous singletons. We estimate θπ (34) using a haploid sample size of 27 for the inbred lines.

### Variant Flanking Sequencing

To identify the most frequent motifs associated with each class of observed single nucleotide changes, we used the Mutation Motif Python package (29). The program performs a comparative statistical analysis of the nucleotide state of positions that immediately flank observed nucleotide variants. The largest effects tend to occur at the positions one or two base pairs up and downstream of variants (29) (see Fig S3). Mutation Motif identifies significant differences in composition at variant sites by drawing a random sample of flanking positions from the immediate context while controlling for positional effects by focusing on a window of sequence around each variant.

We developed a script called SNP_Context (in Bash and Python) that draws samples of variant-adjacent sequence from a reference genome (https://github.com/carte731/SNP_Context). The script reads a VCF file, confirms variant states relative to the reference, checks for overlaps among variant positions, and creates a sequence FASTA file compatible with Mutation Motif (29).

Variant annotation was performed using Variant Effect Predictor (VeP) (37) with gene models provided by (32). VeP reports the total number of transcript changes induced by a variant (Table 1).

**Table 1.** De novo and spontaneous variant consequences, as annotated using Variant Effect Predictor (VEP).

| Consequence (Variant) Type | Number and proportion of affected transcripts |  |  |  |
| --- | --- | --- | --- | --- |
|  | FN | Proportion | Spontaneous | Proportion |
| frameshift | 34 | 0.260 | 17,723 | 0.031 |
| missense | 68 | 0.519 | 292524 | 0.519 |
| synonymous | 29 | 0.221 | 227438 | 0.403 |
| start or stop changes | – | 0.000 | 9517 | 0.017 |
| In-frame indels | – | 0.000 | 14717 | 0.026 |
| other coding | – | 0.000 | 1835 | 0.003 |
| **Total** | **131** |  | **563,754** |  |

To determine the total number of sites in the genome that include the most likely sequence motif for FN mutations, we searched all possible 2-, 3-, 4-mers constituting the most frequent sequence context. We examined sequence context for the two most abundant classes of de novo FN variants. This analysis made use of the EMBOSS (55) compseq tool.

## Acknowledgments

We thank members of the Morrell lab for discussion and software testing. We also would like to thank Li Lei and Shohei Takuno for helpful comments on an earlier version of the manuscript, and Emily Vonderharr for help managing the supplemental data repository. Hardware and software support were provided by the University of Minnesota Supercomputing Institute. This work was supported by the US National Science Foundation Plant Genome Program grant (DBI-1339393 to PLM), the US Department of Agriculture Biotechnology Risk Assessment Research Grants Program (BRAG) (USDA BRAG 2015-06504 to PLM, WAP, SAJ, and RMS).

## Author Contributions

MFR, CKC, RMS, and PLM designed the research. SWR and CKC wrote programs to perform analysis. SRW, MFR, CKC, and PLM performed analysis. SRW, MFR, CKC, WAP, SAJ, RMS, and PLM wrote the manuscript.

---

**Fig S1.**
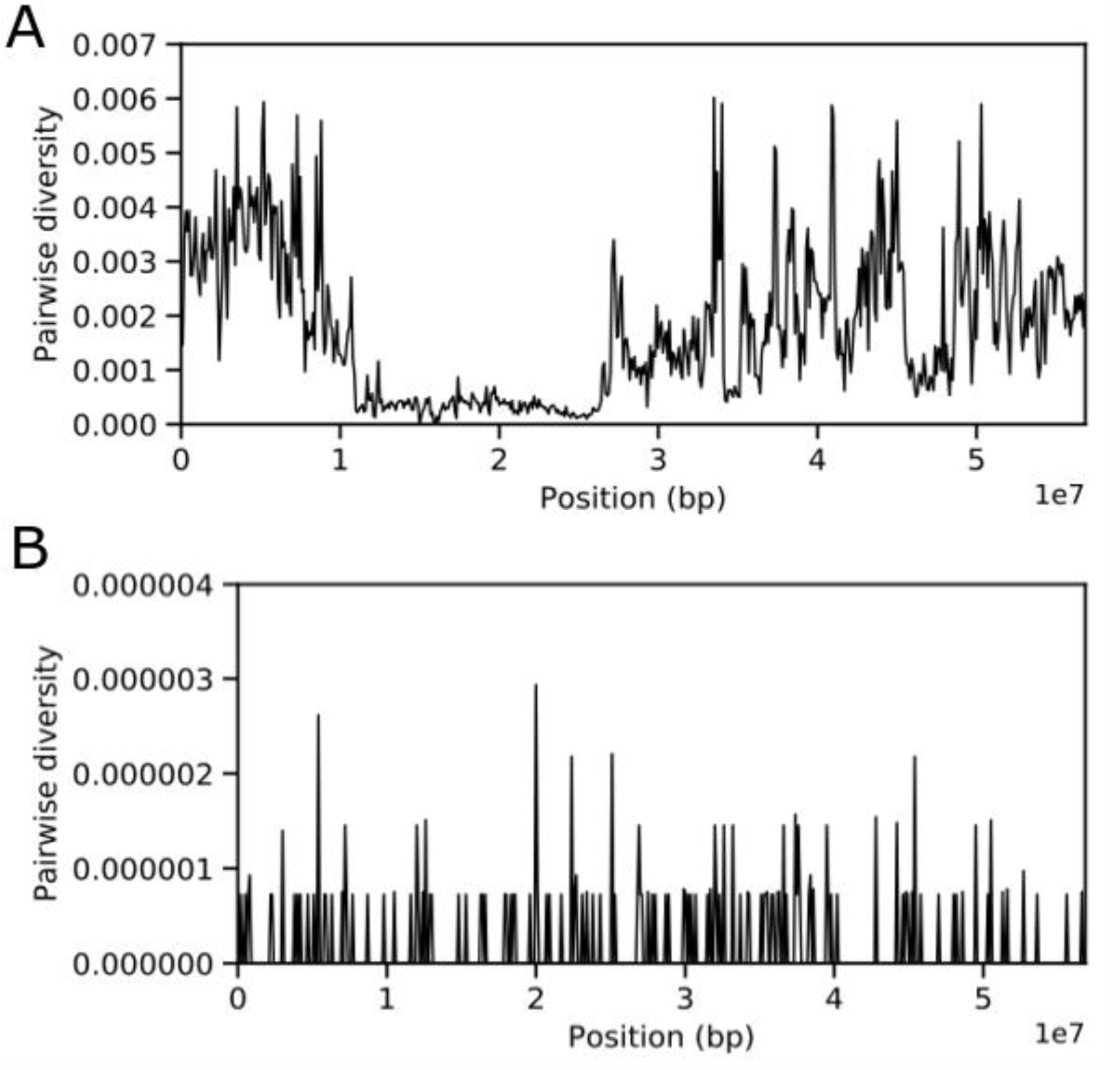
Pairwise diversity in standing variation (panel A) versus FN lines (panel B) across chromosome 1. Note the difference in scale for the y-axis.

**Figure S2.**
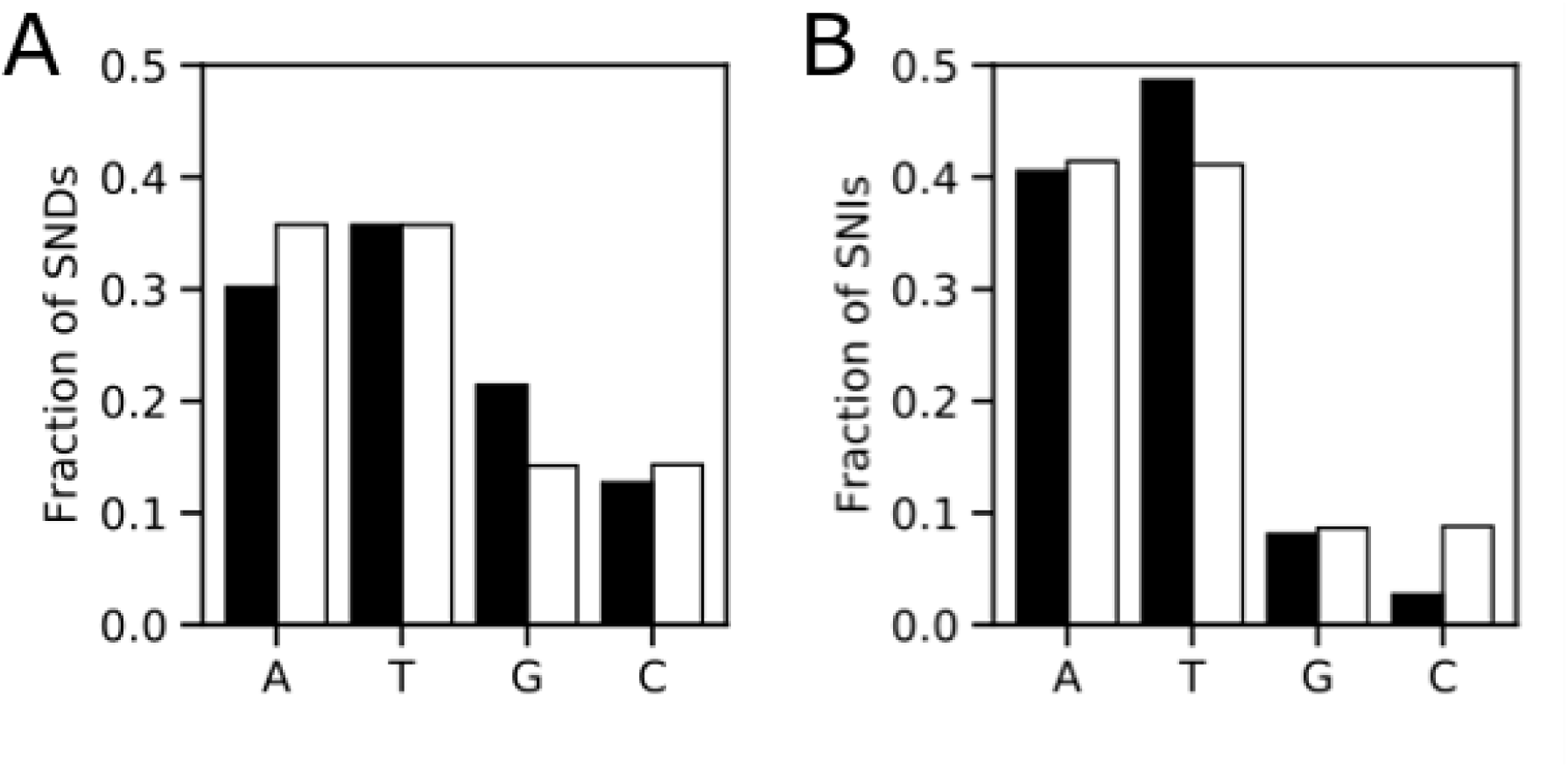
Nucleotide composition of single nucleotide deletions (SNDs) (panel A) and single nucleotide insertions (SNIs) (panel B). Variants in FN lines are shown in black, while variants identified as standing variation are shown in white.

**Figure S3.**
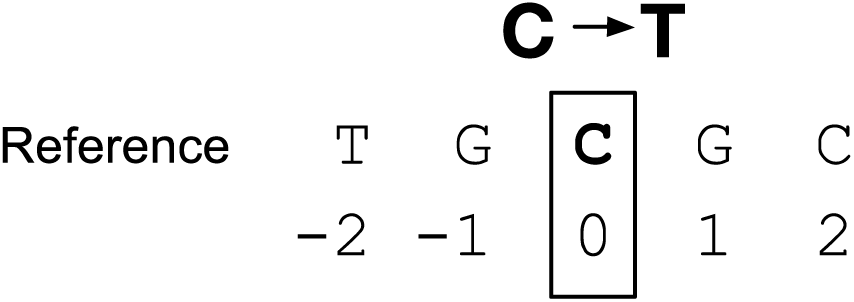
The sequence context for each variant involves the nucleotide sequence in the reference genome that immediately flank a variant nucleotide site. The variant site (in the example, a C->T change) is identified as the “0” position, and positions to the right of the variant identified by positive values and positions to the left identified as negative values.

**Table S1.** Sample information from the fast neutron lines and genetic background line (M92-220.x1.04.WT) from Bolon et al. 2014 that were discarded due to contamination. * Marks three samples which are siblings (family 1) + Marks three samples which are siblings (family 2)

| Sample Name | Generations of Selfing | Fast Neutron Dosage (Gy) |
| --- | --- | --- |
| M92-220.x1.04.WT | N/A | N/A |
| 2012BM7F223 | 7 | 8 |
| 2012CM7F040P05* | 7 | 16 |
| 2012CM7F040P06* | 7 | 16 |
| 2012CM7F040P07* | 7 | 16 |
| 2012CM8F030P02+ | 8 | 16 |
| 2012CM8F030P07+ | 8 | 16 |
| 2012CM8F030P09+ | 8 | 16 |
| 2012DM8F016P02 | 8 | 16 |
| 4R30C22acr626MN13 | 6 | 16 |
| 5R39C03Dr334cMN12 | 3 | 32 |
| FN0112228.06.02.01.M5 | 5 | 16 |
| FN0112885.02.06.03.M5 | 5 | 16 |
| FN0131633.06.01.M4 | 4 | 16 |
| FN0163764.04.01.M4 | 4 | 32 |
| FN0164160.03.02.01.01.M6 | 6 | 32 |
| FN0164472.x3.06.01.M5 | 5 | 32 |
| FN0170228.07.35.01.M5 | 5 | 16 |
| FN0170712.06.41.01.M5 | 5 | 16 |
| FN0171501.01.02.M4 | 4 | 32 |
| FN0172932.09.08.01.M5 | 5 | 16 |
| FN0173217.03.09.01.M5 | 5 | 16 |
| FN0175143.05.06.01.M5 | 5 | 16 |
| FN0175501.x2.02.01.M5 | 5 | 16 |
| FN0190069.01.01.M4 | 4 | 32 |
| R18C55Dhaar437MN13 | 6 | 32 |
| R52C55Dadr564MN13 | 5 | 32 |
| RP8DM5r597MN13 | 6 | 32 |

